# Single-cell analysis of human embryos reveals diverse patterns of aneuploidy and mosaicism

**DOI:** 10.1101/2020.01.06.894287

**Authors:** Margaret R. Starostik, Olukayode A. Sosina, Rajiv C. McCoy

## Abstract

Less than half of human zygotes survive to live birth, primarily due to aneuploidies of meiotic or mitotic origin. Mitotic errors lead to chromosomal mosaicism, defined by multiple cell lineages with distinct chromosome complements. The incidence and fitness consequences of chromosomal mosaicism in human embryos remain controversial, with most previous studies based on bulk DNA assays or comparisons of multiple biopsies of a few embryonic cells. Single-cell genomic data provide an opportunity to quantify mosaicism on an embryo-wide scale. To this end, we extended an approach to infer aneuploidies based on chromosome dosage-associated changes in gene expression by integrating signatures of allelic imbalance. We applied this method to published single-cell RNA sequencing data from 74 disaggregated human embryos, spanning the morula to blastocyst stages. Our analysis revealed widespread mosaic aneuploidies across preimplantation development, with 59 of 74 (80%) embryos harboring at least one aneuploid cell (1% FDR). By clustering copy number calls, we reconstructed histories of chromosome mis-segregation, distinguishing meiotic and early mitotic errors from those occurring after lineage differentiation. We observed no significant enrichment of aneuploid cells in the trophectoderm compared to the inner cell mass, though we do detect such an enrichment in published data from later post-implantation stages. Finally, we observed that aneuploid cells exhibit upregulation of immune response genes, as well as downregulation of genes involved in proliferation, metabolism, and protein processing, consistent with stress responses previously documented in other stages and systems. Together, our work provides a high-resolution view of aneuploidy in preimplantation embryos and supports the conclusion that low-level mosaicism is a common feature of early human development.

## Introduction

Genetic surveys of *in vitro* fertilized (IVF) human embryos consistently reveal substantial levels of aneuploidy—whole chromosome gains and losses that trace their origins to diverse mechanisms of chromosome mis-segregation. These include (primarily maternal) meiotic mechanisms such as non-disjunction, precocious separation of sister chromatids, and reverse segregation (Ottolini et al. 2015), as well as mitotic mechanisms such as mitotic non-disjunction, anaphase lag, and endoreplication (Vázquez-Diez and FitzHarris 2018). In contrast to meiotic errors, which uniformly affect all embryonic cells, mitotic errors generate chromosomal mosaicism, whereby different cells possess distinct chromosome complements. Such mitotic aneuploidies may propagate to descendant cells in a clonal manner and may also contribute to fitness variation. While severe chromosomal mosaicism is lethal to early embryos (McCoy et al. 2015b; Ottolini et al. 2017), low levels of mosaicism appear compatible, and perhaps even common, with live birth (Greco et al. 2015; McCoy 2017).

One major limitation in studying the incidence and implications of chromosomal mosaicism is that most inferences are based on bulk DNA assays or comparisons of multiple biopsies of a few embryonic cells. As a result, current estimates of mosaicism in human embryos range from 4% to 90% (Capalbo et al. 2017). This has provoked intense debate over the true incidence of mosaicism at various developmental stages, its classification as a pathologic versus physiologic state, and its corresponding management in the context of preimplantation genetic testing for aneuploidy (PGT-A) of IVF embryos (Rosenwaks et al. 2018). Specifically, PGT-A seeks to prioritize IVF embryos for transfer based on the ploidy status of embryo biopsies, with current implementations involving biopsies of ~5 trophectoderm cells of day-5 or day-6 blastocysts. This approach is based on the premise that a biopsy is representative of the embryo as a whole and predictive of its developmental outcome. While this premise may be violated by chromosomal mosaicism, the impact of such confounding remains obscure. A more complete picture of aneuploidy across many embryonic cells is therefore critical to a basic understanding of human development, as well as for guiding fertility applications such as PGT-A.

Single-cell genomic datasets offer promising resources for studying mosaic aneuploidy, as they potentially contain valuable information about both cell type and chromosome copy number. Moreover, characteristics of aneuploidies observed in single-cell data may suggest meiotic or mitotic mechanisms of origin. Previous studies have established proof of principle for detecting mosaic aneuploidy using single-cell RNA sequencing (scRNA-seq) data. Griffiths et al. (2017), for example, developed a statistical approach to discover aneuploidies based on chromosome dosage-induced changes in gene expression, validating their method using genome and transcriptome sequencing (G&T-seq) data (Macaulay et al. 2015). Other studies have developed similar approaches for the purpose of studying chromosome instability in cancer (e.g., Fan et al. 2018). In addition to changes in overall expression, aneuploidy is expected to generate allelic imbalance (i.e., allele-specific expression)—deviations from the null 1:1 ratio of expression from maternally and paternally inherited homologs. Here, we extended the expression-based method of Griffiths et al. (2017) to incorporate this complementary signature of allelic imbalance.

Applying this method to scRNA-seq data from 74 embryos (Petropoulos et al. 2016), we quantified the incidence of meiotic and mitotic aneuploidy at single-cell resolution. We observed evidence that both meiotic and mitotic aneuploidies are prevalent across preimplantation stages in patterns consistent with diverse mechanisms of origin. Differences in aneuploidy rates across cell types were not observed in data from preimplantation embryos, but are detectable in published calls from later post-implantation stages. Together, our work provides an embryo-wide census of aneuploidy across early development and quantifies parameters of chromosomal mosaicism that have proven elusive to biopsy-based studies.

## Results

### Detection of aneuploidy in scRNA-seq data

Building upon the foundations of Griffiths et al. (2017; *scploid* software package) we developed an approach to integrate signatures of allelic imbalance to discover aneuploidy in scRNA-seq data (Fig. 1; Methods). Seeking to characterize aneuploidy at single-cell resolution throughout preimplantation development, we applied this approach to published scRNA-seq data from 88 human preimplantation embryos (1529 total cells) spanning the cleavage to late blastocyst stages (E3-E7; Petropoulos et al. 2016). Cell type annotations were obtained from Stirparo et al. (2018) and along with embryonic stage were used to define strata for *scploid*. We removed cells in the lower 10th percentile of mapped reads or percent mapped reads, as well as two cell groups (E3 undifferentiated and E6 epiblast/primitive endoderm intermediate) that failed quality control due to their small numbers of sufficiently expressed genes (Fig. S1). Exclusion of E3 cleavage-stage embryos may also be justified on the basis that the maternal-to-zygotic transition is not yet complete at this stage (Petropoulos et al. 2016). These quality control procedures resulted in the retention of 1115 cells from 74 embryos, spanning the E4 morula to E7 late blastocyst stages (Table S1). Retained stages and cell types exhibited low expression variance, comparable to mouse embryo data used for benchmarking by Griffiths et al. (2017; Fig. S2). Evidence of aneuploidy based on signatures of expression alteration and allelic imbalance was correlated (Fig. S3), but with the latter exhibiting greater sensitivity for detecting monosomy. We combined the two signatures using Fisher’s method (Fisher 1925) to obtain p-values for every cell-chromosome combination (see Methods).

**Fig. 1.**
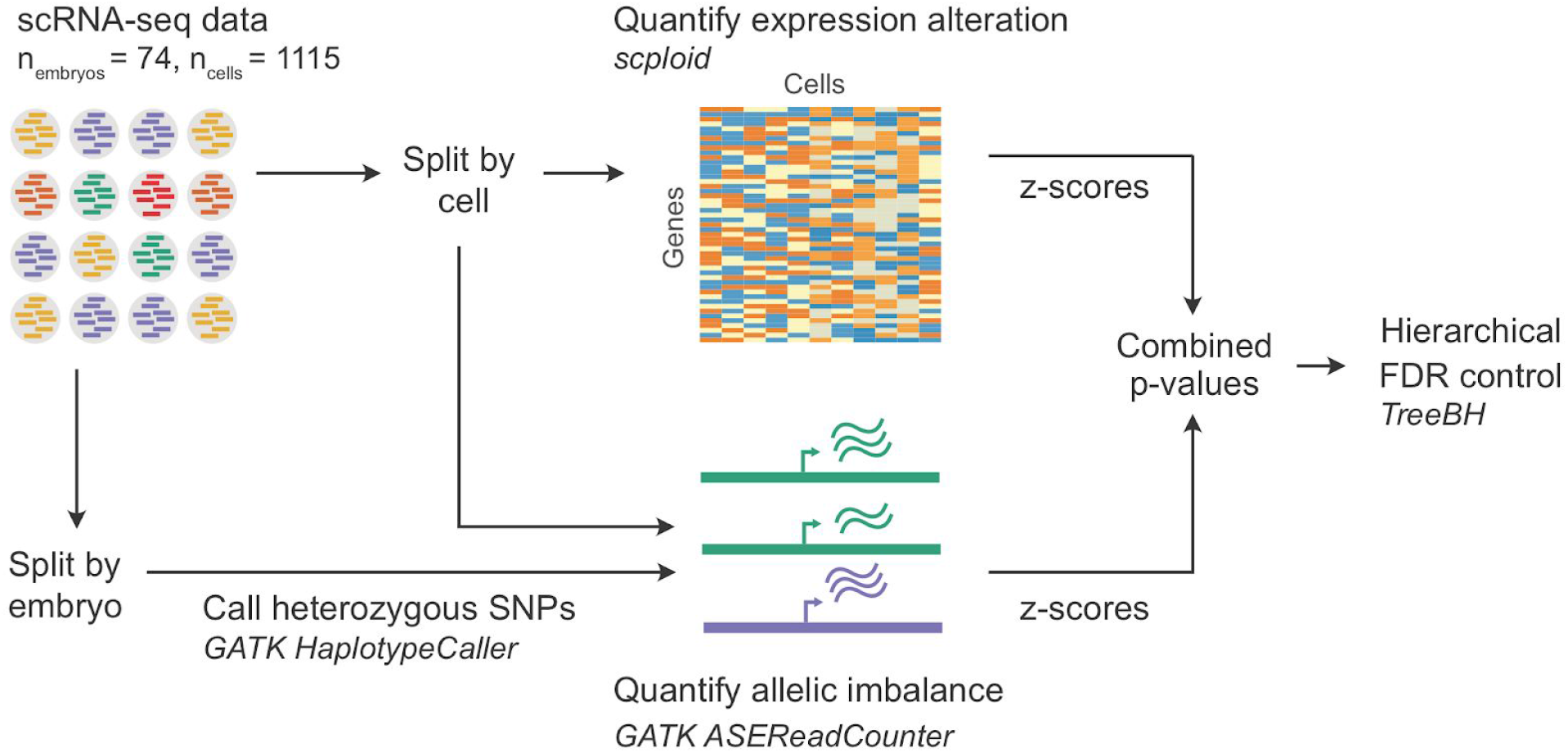
Approach for detecting aneuploidy in single-cell RNA-seq data based on complementary signatures of chromosome-wide gene expression alteration as well as allelic imbalance.

As highlighted in a previous review (Capalbo et al. 2017), the failure to account for multiple hypothesis testing has the potential to inflate estimates of mosaic aneuploidy based on multiple embryo biopsies. This challenge is magnified in single-cell analysis, where each cell-chromosome combination constitutes a separate statistical test (1115 cells · 22 autosomes = 24,530 tests in our study). Meanwhile, answering the most relevant biological questions requires integrating the output of many correlated statistical tests. For example, what proportion of embryos harbor at least one aneuploid cell? What proportion of cells within such embryos are aneuploid? We addressed this challenge using the method *TreeBH* (Bogomolov et al. 2017), an extension of the Benjamini-Hochberg procedure (Benjamini and Hochberg 1995) to tree-structured hypotheses. This allowed us to control the false discovery rate (FDR) at multiple levels (chromosomes, cells, and embryos) while accounting for the hierarchical dependency structure of the data.

Across all cell types and developmental stages, we estimated that 80% (59 of 74) of embryos contained at least one aneuploid cell and that 39% (433 of 1115) of all cells tested across the 74 embryos were aneuploid at an FDR threshold of 1% (Fig. 2A). A total of 4.8% (1172 of 24,530) of all cell-chromosome combinations were called as aneuploid. Patterns of aneuploidy across cells of individual embryos can help distinguish meiotic versus mitotic errors. Because they affect gametes and resulting zygotes, meiotic errors are expected to produce uniform aneuploidies across all embryonic cells. Mitotic aneuploidies meanwhile affect only a fraction of cells, depending upon the timing of their occurrence as well as the possibility of selection against aneuploid cells within mosaic embryos. We found that embryos displayed diverse patterns of aneuploidy, ranging from minor meiotic errors involving one or two chromosomes to chaotic mosaic abnormalities affecting many cells and chromosomes simultaneously (Fig. 2 and Fig. 3). To allow for false negatives, we defined meiotic-origin aneuploidies using 75% as a heuristic cutoff for the percentage of cells per embryo with a particular aneuploidy (i.e., gain or loss of a particular chromosome). All other aneuploidies were classified as mitotic in origin. Based on these criteria, we observed that 5% (4 of 74) embryos possessed only meiotic aneuploidies, 49% (36 of 74) of embryos possessed only mitotic aneuploidies, and 26% (19 of 74) of embryos possessed both meiotic and mitotic aneuploidies.

**Fig. 2.**
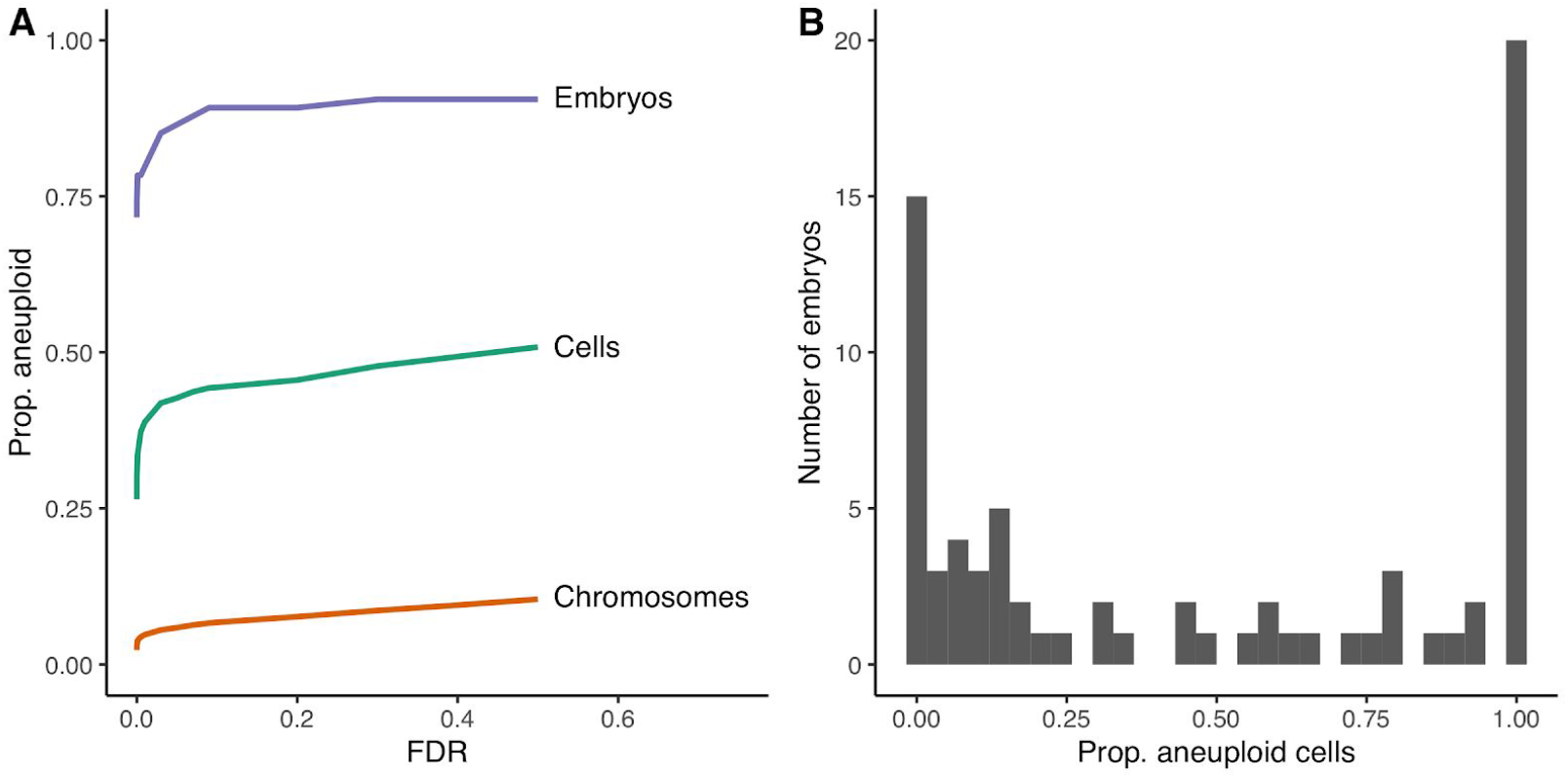
Aneuploidies discovered in scRNA-seq data from human preimplantation embryos (Petropoulos et al. 2016). **A.** Proportions of aneuploid chromosomes, cells, and embryos detected at varying false discovery rates (FDR). Error rates were controlled while accounting for the hierarchical dependency structure of the data (chromosomes within cells within embryos) using TreeBH (Bogomolov et al. 2017). **B.** Distribution of proportions of aneuploid cells per embryo at a 1% FDR.

**Fig. 3.**
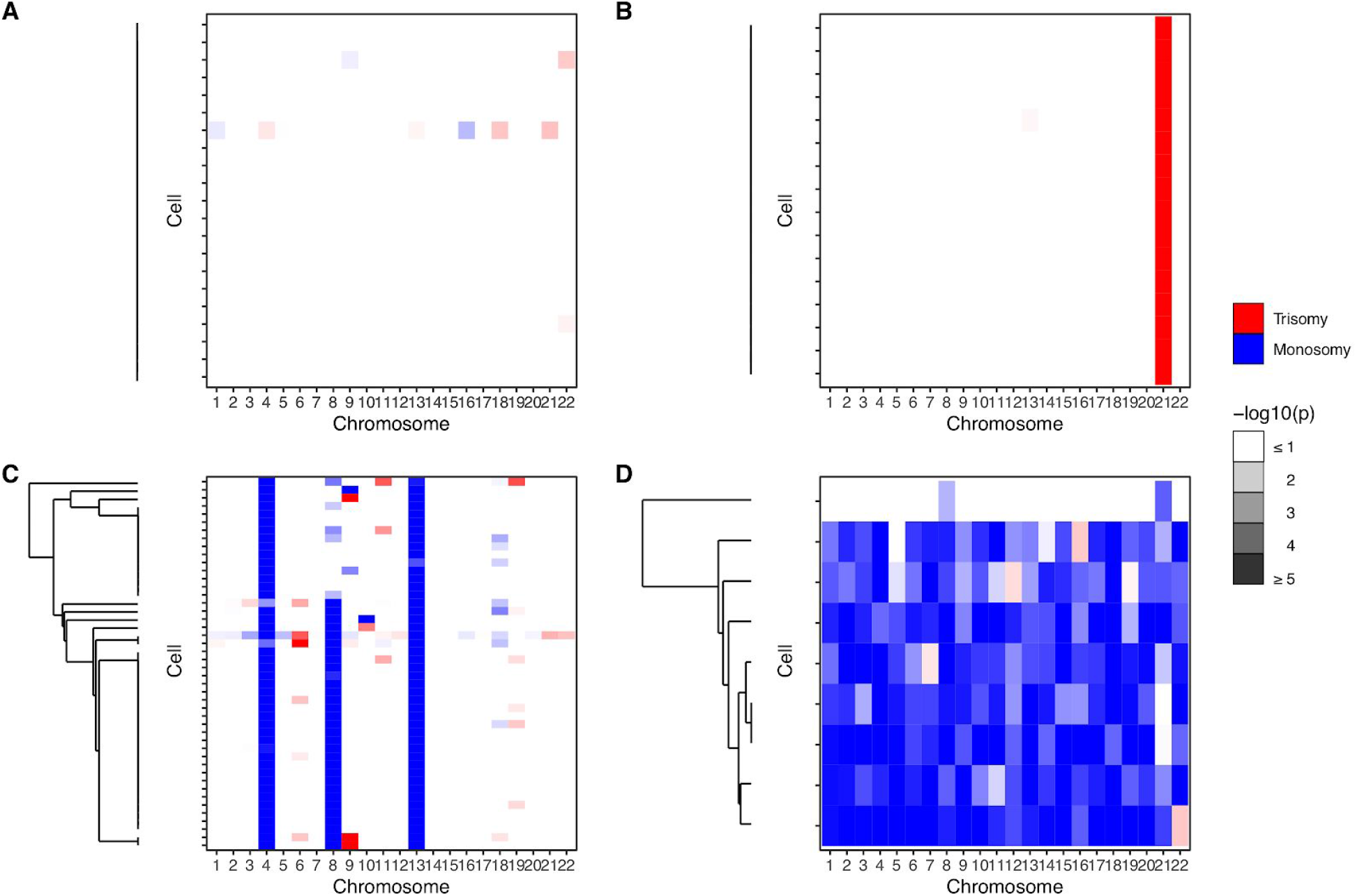
Examples of chromosome abnormalities detected based on scRNA-seq data from human embryos. Each heatmap represents data from an individual embryo. Rows of the heatmaps represent single cells, while columns represent chromosomes (autosomes only). Dendrograms depict hierarchical clustering of aneuploidy signatures, roughly reflecting common ancestry among aneuploid cells. **A.** Embryo E7.3 was called euploid with negligible deviations from the null observed for all chromosomes within all cells. **B.** Embryo E5.13 exhibits a putative meiotic-origin trisomy of chromosome 21. **C.** Embryo E7.17 exhibits putative meiotic-origin monosomies of chromosomes 4 and 13, mosaic monosomy of chromosome 8, and sporadic low-level aneuploidies of other chromosomes. **D.** Embryo E7.5 was inferred as mosaic near-haploid, with haploid or near-haploid signatures in 8 of 9 cells, but near-diploidy in one cell.

The proportion of aneuploid cells per embryo exhibited a characteristic ‘U’-shape, suggesting that meiotic aneuploidies (e.g., Fig. 3B) and low-level mosaic aneuploidies are relatively common, but high-level mosaic aneuploidies are relatively rare (Fig. 2B). Blastocyst E7.17 provides an intriguing example of multiple forms of aneuploidy within a single embryo (Fig. 3C). Specifically, chromosomes 4 and 13 displayed evidence of meiotic origin monosomy, while chromosome 8 displayed evidence of mosaic monosomy affecting approximately half of cells. The latter observation is potentially consistent with chromosome loss (e.g., via anaphase lag) during the first embryonic cleavage. Meanwhile, other chromosomes of this embryo displayed evidence of sporadic low-level aneuploidy, such as monosomy of chromosome 10. An even more extreme form of mosaicism was detected in blastocyst E7.5, which we inferred to be mosaic near haploid (Fig. 3D). Seven of the eight cells showed chromosome-wide monoallelic expression, while one cell showed mostly biallelic expression. Due to its severe nature, this abnormality was only detectable based on signatures of allelic imbalance, as analysis based on overall expression alteration lacked a baseline for comparison (Fig. S4).

While we observed a negative correlation between chromosome-specific aneuploidy rates and the number of protein-coding genes per chromosome (Pearson’s r = −0.546, p = 8.64 × 10^−3^; Fig. S5), differences in aneuploidy rates among chromosomes were not significant upon accounting for non-independence among chromosomes within cells within embryos (χ^2^(df = 21, n = 24,530) = 29.0, p = 0.114; see Methods).

### Cell-type specific variation in aneuploidy may arise and intensify during postimplantation development

Long-standing questions in the field of preimplantation genetics include how aneuploid cells are distributed among different cell types and how this changes throughout development. Cell type-specific propensities and/or tolerances for aneuploidy could help explain observations such as confined placental mosaicism observed at later developmental stages (Toutain et al. 2018). Meanwhile, selection against aneuploid cells within mosaic embryos could help explain recent reports that some embryos that test mosaic with PGT-A can result in healthy live births after intrauterine transfer (Greco et al. 2015). Bolton et al. (2016) previously used a chimeric mouse model to address these questions, demonstrating that aneuploid cells of the inner cell mass undergo apoptosis, while aneuploid cells of the trophectoderm are tolerated but experience proliferative defects. While groundbreaking, the relevance to human development has remained uncertain, as mouse embryos are known to exhibit lower rates of chromosome instability and higher rates of blastocyst formation than their human counterparts (Daughtry and Chavez 2016). Investigating these processes in human embryos is therefore essential for understanding the cellular and organismal fitness consequences of chromosomal mosaicism.

We obtained cell type annotations of the Petropoulos et al. (2016) dataset from Stirparo et al. (2018) and confirmed that these groups formed clusters based on uniform manifold approximation and projection (UMAP) dimension reduction of the gene expression matrix (Fig. 4A,B). We then examined the relationship between aneuploidy status and cell type using a binomial generalized linear mixed model (GLMM) with embryo as a random effect and embryonic stage (days post-fertilization) and cell type annotation as fixed effects (see Methods). We estimated the average marginal effect (AME) for a given predictor variable as its effect per cell, averaged across cells (see Methods). We compared this model to a reduced model without the cell type term to test whether aneuploidy rates varied across cell types. We detected no significant difference in aneuploidy rates across cell types (χ^2^(df = 5, n = 1115) = 6.19, p = 0.288; Figure 4F) or across days of development (E4 to E7; AME = −0.071, SE = 0.046, p = 0.124). We similarly detected no significant enrichment of aneuploidy in the trophectoderm versus the inner cell mass and its descendant lineages (AME = 0.012, SE = 0.039, p = 0.765). We note, however, that the wide confidence interval (95% CI [−0.064, 0.087]) signifies that we cannot rule out modest differences.

**Fig. 4.**
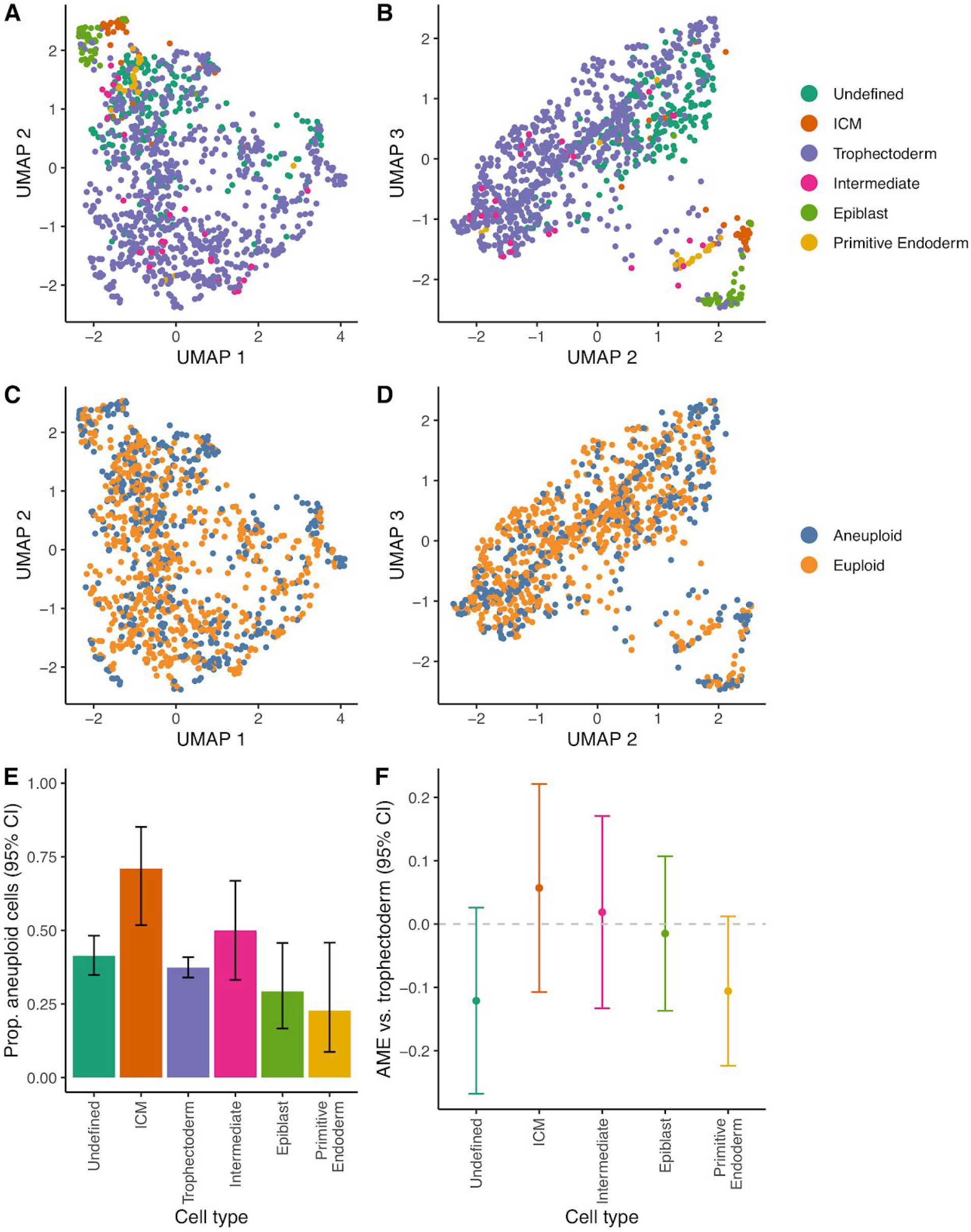
Comparisons of aneuploidy across cell types. **A.** Individual cells plotted on the first and second UMAP dimensions, colored by cell type annotations from Stirparo et al. (2018). **B.** Same as panel ‘A’, but for the second and third UMAP dimensions. **C.** Cells plotted on the first and second UMAP dimensions, colored by aneuploidy status. **D.** Same as panel ‘C’, but for the second and third UMAP dimensions. **E.** Proportions of aneuploid cells, stratified by cell type. **F.** Average marginal effects (AME) of cell types on aneuploidy rates relative to aneuploidy rates of trophectoderm cells—the source for PGT-A biopsies. Confidence intervals of all estimates overlap zero, indicating no significant difference for any cell type.

**Fig. 5.**
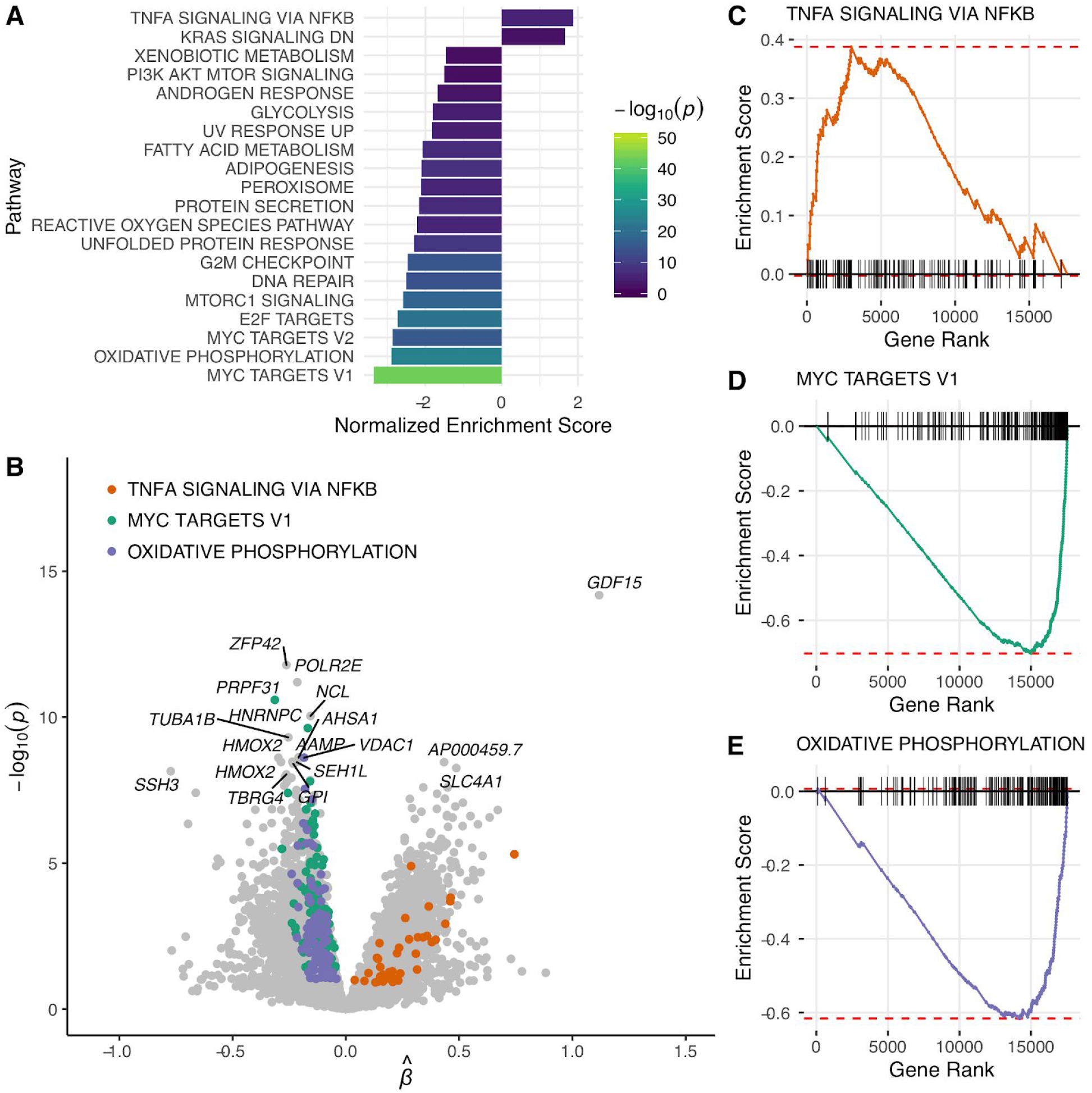
Transcriptional responses to aneuploidy in human embryos. **A.** Hallmark gene sets from the Molecular Signatures Database (MSigDB) that are significantly enriched for genes that are up-or down-regulated in aneuploid cells based on gene set enrichment analysis (GSEA; 5% FDR). **B.** Volcano plot depicting differential expression between euploid and aneuploid cells. Positive values indicate increased expression in aneuploid cells, while negative values indicate reduced expression. **C.** Gene set enrichment plot demonstrating that genes that genes regulated by NF-kB in response to tumor necrosis factor-α are significantly up-regulated in aneuploid cells. **D.** Same as panel ‘C’, but demonstrating that Myc targets exhibit reduced expression in aneuploid cells. **E.** Same as panel ‘C’, but demonstrating that genes involved in oxidative phosphorylation are downregulated in aneuploid cells.

Though informative of cell type, the sparse and bursty nature of scRNA-seq data poses a challenge for aneuploidy inference, placing practical limits on sensitivity and specificity (Griffiths et al. 2017). We thus sought to replicate the qualitative patterns of mosaic aneuploidy that we previously described using published data from additional disaggregated embryos that were analyzed by single-cell post-bisulfite adaptor tagging (PBAT) DNA methylome sequencing (Zhu et al. 2018). The fact that these data were based on single-cell DNA-sequencing (scDNA-seq) lends confidence to the aneuploidy calls. To facilitate comparison with our scRNA-seq results above, we focused on the twenty embryos from the morula and blastocyst stages of development. A total of 65% (13 of 20) of these embryos possessed at least one cell called as aneuploid. Applying the same definition we previously described (75% of cells of an embryo possessing a particular monosomy or trisomy), 9 (45%) of these embryos possessed only mitotic aneuploidies, one (5%) embryo possessed only meiotic aneuploidies, and 3 (15%) embryos possessed both meiotic and mitotic aneuploidies. Hierarchical clustering of these aneuploidy calls revealed patterns qualitatively consistent with our scRNA-based results, including prevalent low-level mosaicism (Fig. S6). Among the twelve blastocyst-stage embryos that could be tested, we detected no significant difference in the rates of aneuploidy between cells of the trophectoderm versus the inner cell mass (AME = 0.014, SE = 0.058, p = 0.811), again consistent with our scRNA-seq based results.

A recent study by Zhou et al. (2019) developed an extended *in vitro* culture system to produce the first single-cell genomic data from postimplantation human embryos spanning days 6 to 14 of development. This included Trio-seq data (including single-cell bisulfite sequencing) from 17 embryos, as well as scRNA-seq data from an additional 48 embryos. Applying the GLMM described above to published single-cell aneuploidy calls from these postimplantation embryos, we detected a significant enrichment of aneuploid cells in the trophectoderm compared to the epiblast and primitive endoderm (lineages derived from the inner cell mass). This enrichment was detected in scDNA-based calls from 286 cells of the 17 bisulfite-sequenced embryos (AME = 0.142, SE = 0.064, p = 0.028), as well as scRNA-based calls from 5911 cells of the 48 additional embryos (AME = 0.049, SE = 0.022, p = 0.024). Intriguingly, the latter sample also revealed that the enrichment of aneuploid cells in the trophectoderm became stronger throughout post-implantation development (β_lineage × stage_ = 0.212, SE = 0.068, p = 1.84 × 10^−3^). We note that this interaction model includes a random effect of embryo, thus addressing correlations among cells by allowing embryos to vary in their baseline rates of aneuploidy.

### Global gene expression responses to aneuploidy

In addition to the primary (i.e., *cis*-acting) and secondary (i.e., *trans*-acting) dosage effects, aneuploidy may induce tertiary transcriptional changes, including responses to proteotoxic, oxidative, and hypo-osmotic stresses (Dürrbaum et al. 2014; Tsai et al. 2019). To investigate this phenomenon in the context of human preimplantation development, we used a negative binomial mixed model to test for differential expression between euploid and aneuploid cells (see Methods). Embryo and cell type were specified as random effects to again account for the correlation among cells within embryos, while embryonic stage (days post-fertilization) and aneuploidy status were specified as fixed effects (see Methods).

Using this model, we identified 2925 genes that were differentially expressed between euploid and aneuploid cells (5% FDR; Table S2). The most significant association involved upregulation of the *Growth/differentiation factor 15 (GDF15)* in aneuploid relative to euploid cells (β = 1.118, SE = 0.144, p = 6.6 × 10^−15^; Fig. S7). Intriguingly, this gene was previously discovered to be upregulated in aneuploid human cell lines compared to diploid cell lines from which they were derived, suggesting that *GDF15* may serve as a biomarker of aneuploidy across stages and cell types (Dürrbaum et al. 2014). The gene *ZFP42* exhibited the most significant downregulation in aneuploid versus euploid cells (β = −0.262, SE = 0.037, p = 1.6 × 10^−12^; Fig. S8). This gene encodes the zinc finger protein REX1—a classic marker of pluripotency whose expression contributes to lineage specification during early development (Son et al. 2013). The role of aneuploidy in altering such lineage decisions via *ZFP42* downregulation may therefore merit future investigation.

To gain further insight into global responses to aneuploidy in human embryos, we performed gene set enrichment analysis on the hallmark gene sets from the Molecular Signatures Database (MSigDB). This analysis revealed 20 gene sets that were significantly enriched among the tails of differentially expressed genes (5% FDR). Notably, we observed fewer gene sets significantly enriched for genes that are up-(2 gene sets) versus down-regulated (18 gene sets) in aneuploid cells. Downregulated gene sets included those related to cell proliferation, protein processing, and metabolism. Gene sets enriched for genes upregulated in aneuploid cells included those that are downregulated by KRAS signaling (again potentially reflecting an anti-proliferation response) as well as genes regulated by NF-κB in response to TNF-α, broadly consistent with cellular stress and inflammatory signals previously reported in aneuploid cell lines (Santaguida et al. 2017; Liu et al. 2017).

## Discussion

One key limitation of most previous studies of aneuploidy in human preimplantation embryos has been their reliance on biopsies of one or few cells. Many studies have adopted an operational definition of mosaicism based on PGT-A results that are intermediate between those expected of uniform euploid and uniform aneuploid biopsies (Cram et al. 2019). This narrow definition of mosaicism ignores the possibility of aneuploidy among the non-biopsied cells that compose the rest of the embryo. In contrast, a biological definition of mosaicism denotes the presence of cells with distinct chromosome complements anywhere within the embryo. While mathematical modeling approaches (e.g., Gleicher et al. 2017) can help reconcile studies based on disparate definitions, such models require assumptions about unknown parameters including the spatial and lineage-specific distributions of aneuploid cells within mosaic embryos.

We sought to overcome these limitations by leveraging published scRNA-seq data from disaggregated human embryos. By combining signatures of gene expression alteration and allelic imbalance, we revealed patterns of meiotic and mitotic aneuploidy at single-cell resolution. A total of 31% of embryos displayed uniform or near-uniform aneuploidy of at least one chromosome across all cells—a pattern attributable to meiotic errors, which largely trace to maternal oogenesis. Meanwhile, low-level mosaicism was prevalent across all cell types and developmental stages, with 74% of embryos inferred to possess at least one cell affected by mitotic error. While substantially higher than most biopsy-based studies, this estimate is roughly in line with the few previous studies to quantify aneuploidy in single cells of disaggregated embryos, albeit with different methodologies or at different stages. A landmark study by Vanneste et al. (2009) used SNP genotyping and array comparative genomic hybridization (CGH) to analyze 86 single cells from 23 disaggregated cleavage-stage embryos, finding that only three embryos were uniformly diploid, while the rest were either diploid-aneuploid mosaics or mosaics of entirely aneuploid cells. A recent study of 49 disaggregated cleavage-stage rhesus macaque embryos used scDNA-seq to demonstrate that 13 were euploid, 9 were affected by solely meiotic errors, and the remaining 27 by mitotic errors or errors of ambiguous origin (Daughtry et al. 2019). Our estimates are also roughly consistent with aneuploidy calls based on scDNA-and scRNA-seq of human blastocysts (Zhu et al. 2018; Zhou et al. 2019), which reported evidence of mitotic-origin aneuploidy in more than half of embryos.

One surprising observation from our study was the discovery of a mosaic near-haploid embryo (embryo E7.5) in which 8 of 9 cells appeared haploid or near-haploid, but one cell appeared near-diploid. This extreme form of mosaicism escaped detection based on gene expression analysis alone, but was evident based on signatures of allelic imbalance. Hydatidiform moles are known to affect approximately 1 in 600 pregnancies, half of which are triploid dispermic (two paternal and one maternal set of chromosomes) and half of which are diploid androgenetic (two paternal sets of chromosomes). An estimated 85% of the latter type are monospermic, and may arise via the extrusion of maternal chromosomes to the first polar body, followed by “diploidization” of the paternal chromosomes (Nguyen et al. 2018). The mosaic near-haploid constitution of embryo E7.5 is theoretically consistent with a dispermic origin, as a result of postzygotic diploidization of a triploid zygote (Golubovsky 2003). Uniparental diploidy is indistinguishable from haploidy with our approach. Fertilization with two sperm would explain the biallelic nature of the diploid cell, while the dispermic transmission of supernumerary centrioles could also induce mosaicism via multipolar mitosis. While the IVF procedures used to produce the embryos sequenced by Petropoulos et al. (2016) were not reported, the growing use of intracytoplasmic sperm injection (ICSI) has dramatically reduced the prevalence of dispermy. Meanwhile, work in bovine embryos has revealed that even normally fertilized zygotes may produce mixoploid embryos by a mechanism termed “heterogoneic division” (Destouni et al. 2016). Specifically, chromosomes may segregate on an atypical gonomeric spindle to produce a mixture of androgenetic, gynogenetic, and normal diploid daughter cells, thus providing one alternative mechanism to explain embryo E7.5. We anticipate that future large-scale studies of disaggregated human embryos will reveal novel forms of mosaicism whose mechanisms of origin remain to be described.

By leveraging cell type information contained within scRNA-seq data, we also evaluated the long-standing question of how aneuploidy rates vary among cell types. This question is intriguing from both basic biological and clinical perspectives. Such comparisons provide insight into the developmental landscape of gene essentiality and dosage sensitivity, while also shedding light on the representativeness of PGT-A biopsies obtained from trophectoderm tissue. We detected no significant enrichment of aneuploidy within the trophectoderm, but note that the wide confidence interval (AME = 0.012, 95% CI [-0.064, 0.087]) indicates that we cannot rule out modest differences. Indeed, the ~6% enrichment of aneuploidy observed in trophectoderm cells of mature mouse blastocysts by Bolton et al. (2016) falls within this interval. Nevertheless, our study places bounds on any potential differences and provides a useful quantitative framework for testing such hypotheses in future single-cell datasets. While we replicated this lack of enrichment using scDNA-seq-based calls from additional preimplantation embryos (Zhu et al. 2018), we detected significant enrichment of aneuploidy in the trophectoderm versus the primitive endoderm and epiblast (lineages derived from the inner cell mass) in published data from postimplantation embryos (Zhou et al. 2019). Notably, the latter dataset included nearly 6000 cells, lending greater statistical power to such comparisons. The aneuploidy calls from Zhou et al. (2019) also revealed a significant interaction with day of development, indicating that the enrichment of aneuploidy in the trophectoderm becomes more extreme as development proceeds. Whether this observation reflects cell-type-specific apoptosis and/or proliferation defects of aneuploid cells merits future investigation.

The transcriptional consequences of aneuploidy are known to extend beyond direct dosage effects to *trans*-acting impacts on other chromosomes as well as tertiary stress responses (Sheltzer et al. 2012; FitzPatrick 2005). Several recent studies have examined these responses using bulk RNA-seq analysis of embryos with specific aneuploidies (Sanchez-Ribas et al. 2019; Weizman et al. 2019; Kawai et al. 2018; Licciardi et al. 2018). However, bulk RNA-seq averages effects across cells, and studies of specific aneuploidies may conflate primary, secondary, and tertiary effects, thus hindering interpretation. We used a negative binomial mixed effects model to examine indirect responses to aneuploidy in single cells throughout early human development. This statistical method effectively accounts for non-independence among cells within embryos. Such sampling designs are common to many scRNA-seq datasets, and mixed effects models may be broadly applicable to differential expression analysis in this context. Our analysis revealed thousands of genes that are differentially expressed in aneuploid versus euploid cells. The top associated gene was a known biomarker of aneuploidy (*GDF15;* Dürrbaum et al. 2014), thus supporting our approach. While statistically robust, the distributions of *GDF15* expression in euploid and aneuploid cells overlapped substantially (Fig. S7), underscoring the conclusion that no individual gene is diagnostic of aneuploidy status. Nevertheless, our results may be useful for exploring signatures of aneuploidy and associated stress responses, which could in turn be correlated with developmental outcomes. Indeed, multiple previous studies have suggested the utility of gene expression signatures for IVF embryo selection, but have yet to be validated in large independent datasets (Vera-Rodriguez et al. 2015; Groff et al. 2019).

A key methodological advance described here was the integration of signatures of expression alteration and allelic imbalance to detect aneuploidy in single cells. Consideration of allelic imbalance bolsters confidence in our results, especially in cases of monosomies, which generate monoallelic expression across entire chromosomes (allowing for technical artifacts such as barcode swapping, spurious SNPs, and mis-mapped reads). Despite this advance, inference of aneuploidy from scRNA-seq data remains challenging. An important caveat is that any deviation of expression and allelic balance from diploid expectations could lead us to reject the null hypothesis, while phenomena other than whole-chromosome aneuploidy may occasionally induce such deviations. For example, large structural variation may be falsely classified as aneuploidy by our approach. While the effect size thresholds that we implemented help mitigate this concern, approaches to explicitly distinguish segmental and whole-chromosome aneuploidies are an important area for future development. Additional opportunities for methodological improvement include the integration of population-based and/or read-based phasing into allelic imbalance analysis, which could be especially beneficial for the inference of trisomies (Loh et al. 2018).

One additional caveat is that the data analyzed in our study derive from IVF embryos obtained from relatively few patients, about whom no demographic or clinical information was published. We therefore urge caution in extrapolating these findings to a broader population. Previous studies have established a strong association between maternal age and incidence of meiotic error in preimplantation embryos (Hassold and Hunt 2001). Studies have also revealed significant, albeit modest associations between aneuploidy rates and various fertility diagnoses (McCoy et al. 2015b; Kort et al. 2018), as well as patient genotypes (McCoy et al. 2015a; Chernus et al. 2019). One persistent concern with all studies of preimplantation embryos is the possibility that IVF culture conditions impact chromosome stability. Indeed, such impacts have been documented by comparing *in vitro* versus *in vivo* matured bovine embryos (Tšuiko et al. 2017), though no such differences have been detected in humans during preimplantation development (Munné et al.) or at live birth (Zamani Esteki et al. 2019).

Aneuploidy is the leading cause of pregnancy loss and congenital birth defects in humans (Hassold and Hunt 2001). As genetic testing platforms have improved, the existence of chromosomal mosaicism is increasingly acknowledged, but the prevalence of this phenomenon remains disputed and the impacts on human development remain unclear (McCoy 2017). Here we developed an approach to leverage scRNA-seq data from disaggregated human embryos to quantify aneuploidy and mosaicism at single-cell resolution. Our results support the conclusion that meiotic aneuploidies and low-level mosaic aneuploidies are common, but high-level mosaic aneuploidies are relatively rare. Aneuploidy rates among various cell types are similar during preimplantation development, but may arise and intensify throughout postimplantation development. Together, our study reconciles disparate estimates of mosaicism based on different definitions and provides a quantitative framework for investigating aneuploidy in ever-growing single cell datasets.

## Methods

### Aneuploidy inference on scRNA-seq data

Aneuploidy inference was based on complementary signatures of chromosome-wide differential expression and allelic imbalance. The differential expression signature is the basis of the software package *scploid*, which was previously benchmarked using genome and transcriptome sequencing (G&T-seq) data from mosaic aneuploid mouse embryos.

Gene expression quantifications from the Petropoulos et al. (2016) dataset were obtained from conquer (Soneson and Robinson 2018) as input to *scploid*. Analysis was limited to the autosomes, as required by the software. Cells in the lower 0.1 quantile of mapped reads and/or percent mapped reads were excluded from analysis. Cell type annotations were obtained from Stirparo et al. (2018) and visualized on both principal component (Fig. S1A) and UMAP (Fig. 4A-B) dimensions using Monocle (Trapnell et al. 2014) to confirm their clustering. Embryonic stage and cell type annotations were used to define strata as input to *scploid*, thereby limiting cell type-and stage-specific variation that may confound aneuploidy inference. Groups that failed *scploid* quality control procedures were dropped from the analysis (Fig. S1B). This resulted in the inclusion of 1,115 cells from 74 embryos across eleven stage/cell type groups. Each group expressed between 3053 and 3351 genes at a median of 50 counts per million reads mapped (CPM) or greater per cell.

### Allelic imbalance z-scores

Raw single-cell RNA-seq data from Petropoulos et al. (2016) were obtained from EMBL-EBI Array Express (E-MTAB-3929). Reads were mapped to the reference genome using STAR (v2.7.1a). Single-cell alignments from the same embryo were then merged using samtools (v1.9), and processed for variant discovery with GATK (v4.0.12.0), according to the following workflow: https://gatkforums.broadinstitute.org/gatk/discussion/3891/calling-variants-in-rnaseq. Using embryo-level heterozygous SNPs and corresponding single-cell alignments as input, allelic read counts were then computed at every heterozygous SNP in every cell using the ASEReadCounter tool of GATK. The minimum of the counts of reference and alternative allele-supporting reads was obtained for every heterozygous SNP, summed across the chromosome, then divided by the total read count for that chromosome to obtain an allelic imbalance ratio.

To convert the observed allelic imbalance ratios to z-scores, we first regressed out the effect of the number of reads on allelic imbalance since we observed a positive correlation between the observed proportions and the number of reads used to generate them. Then, under the assumption that most cell-chromosome combinations would lie under the null, we identified all null data points as those whose allelic imbalance estimates lie between the first quartile and the third quartile of the empirical distribution. We then estimated the residual at the null proportion as the mean of all the generated residuals for the null data points. To estimate the variance under the null, we further assumed that the residuals were approximately normally distributed. With this assumption and recalling that we already assumed that most cell-chromosome combinations would lie under the null, we derived the variance of the residual of the allelic imbalance under the null using the formula below,

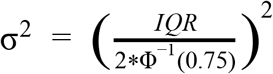

where IQR is the empirical interquartile range of the residuals, and Φ^−1^ is the inverse of the standard normal cumulative distribution function. This is motivated by the fact that under the normal distribution, the interval, I, contains the middle 50% of the data as

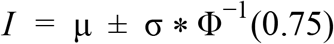

This interval has length,

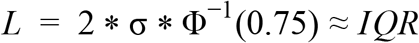

Hence, solving for σ and squaring it, we obtain the result above. With the mean and variance obtained under the null, we then converted the residuals to z-scores by subtracting this mean from each point and dividing by the square root of the variance. We then computed one-sided p-values, based on the expectation that both monosomy and trisomy will have the effect of increasing allelic imbalance.

### Omnibus test combining allelic imbalance and sciploid results

We conducted an omnibus test for each cell-chromosome using the p-values obtained from the scploid and allelic imbalance analyses described above. We note that as a last step, *scploid* imposes an effect size threshold (|1 - *s_ij_*| ≥ 0.2) to classify a chromosome as aneuploid (Griffiths et al. 2017). To incorporate this threshold into our analysis, we set *scploid* p-values for cell-chromosomes below this threshold to 1. We combined allelic imbalance and *scploid* p-values using Fisher’s method (Fisher 1925). Correction for multiple testing was then carried out using *TreeBH* (Bogomolov et al. 2017) to account for the hierarchical nature of the data. Hypothesis tree structure was defined as chromosomes nested within cells nested within embryos, and FDR was controlled at 1% at each level. To assign aneuploid chromosomes to the categories of monosomy and trisomy, we applied k-means clustering (*k* = 2) to the z-scores from the allelic imbalance and *scploid* analyses. Cell-chromosomes composing the cluster with lower mean z-scores were classified as monosomic, and cell-chromosomes composing the other cluster were classified as trisomic.

### Cell-type specific propensity for aneuploidy

We assessed the effect of cell-type on aneuploidy status using mixed-effects logistic regression. We fit two models. In both models, our outcome was a binary variable indicating aneuploidy status (defined as aneuploid if the cell possessed one or more aneuploid chromosome) and embryonic stage (days post-fertilization), which was treated as a fixed effect continuous variable. In the first model, we included cell-type (categorical variable), and in the second model, we included an indicator variable for the trophectoderm cells (with all other cell types grouped as non-trophectoderm). We treating embryo as a random effect to account for correlation among cells within embryos and estimated the random intercept for both models. Embryos were meanwhile assumed independent of one another.

### Average Marginal Effects (AME)

Let E(*Y_ij_*| *X_t_*, *b_j_*) represent the fitted values from each of the generalized linear mixed models defined above for the ith cell with the jth random effect (for example, in the cell-type specific propensity analysis, j refers to the jth embryo) and X is the covariate whose AME we wish to calculate. The AME is defined for a binary covariate X as

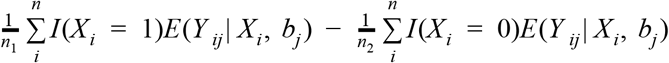

Where 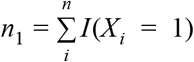 and 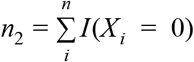. We estimate these values and their standard errors using the *margins* package in R (Leeper et al. 2018).

### Differential expression and gene set enrichment analysis

Global transcriptional responses to aneuploidy were investigated by testing for differential expression between cells called as euploid versus aneuploid. Single cell expression counts were normalized using *SCnorm* (Bacher et al. 2017), with cell type annotations specified as biological conditions. Analysis was limited to broadly expressed genes with ≥1 normalized expression count for at least half of cells. To mitigate direct dosage (i.e., *cis*-acting) effects of aneuploidy, we restricted each test to cells called as euploid for the chromosome containing the respective gene. For each gene, we then fit a negative binomial mixed model, implemented with *lme4* (Bates et al. 2014, 4). Normalized read counts (plus a pseudocount) were specified as the response variable, cell type and embryo were specified as crossed random effects, and embryonic stage (i.e., day post-fertilization) and aneuploidy status were specified as fixed effects. We fit both a random slope and intercept, as well as a random intercept-only model for the cell type variable, retaining the more complex model only if it significantly improved fit over the reduced model based on analysis of deviance (α = 0.05). Coefficients, test statistics, and p-values were evaluated for the aneuploidy status term. Models producing convergence warnings (2.1% or 383 of 17,970 genes) were dropped from the analysis.

Gene set enrichment analysis (GSEA; Subramanian et al. 2005) was performed using the *fgsea* (Korotkevich et al. 2019) package in R. In order to limit the number of tests and improve biological interpretability, we focused our analysis on the Molecular Signatures Database (MSigDB) hallmark gene sets (Liberzon et al. 2015), accessed via *msigdbr* (Dolgalev 2019). Genes were ranked by signed p-value as input to GSEA, which was run using the adaptive multilevel splitting option to compute arbitrarily small p-values.

## Data access

Raw single-cell RNA-seq data from Petropoulos et al. (2016) were obtained from EMBL-EBI Array Express (E-MTAB-3929), while processed gene expression quantifications were obtained from conquer (Soneson and Robinson 2018). Aneuploidy calls based on single-cell DNA and RNA-sequencing of additional embryos were obtained from supplementary tables of Zhu et al.

(2018) and Zhou et al. (2019), while raw data were obtained from GEO (Accession: GSE81233). All code necessary for reproducing our analyses is available at https://github.com/mccoy-lab/aneuploidy scrnaseq (doi:10.5281/zenodo.3599806).

## Acknowledgments

Thank you to staff at the Maryland Advanced Research Computing Center for computing support. Thank you also to the Origins of Aneuploidy Research Consortium for critical feedback. Thanks to Alexis Battle, Rong Li, Jin Zhu, as well as members of the McCoy lab for helpful discussion. This work is supported by NIH grant R35GM133747 to R.C.M.

## Disclosure declaration

The authors declare no competing interests.

## Notes

https://github.com/mccoy-lab/aneuploidy_scrnaseq

