## Supplementary Figures and Tables for "Single-cell analysis of human embryos reveals diverse patterns of aneuploidy and mosaicism"

### Supplementary Information

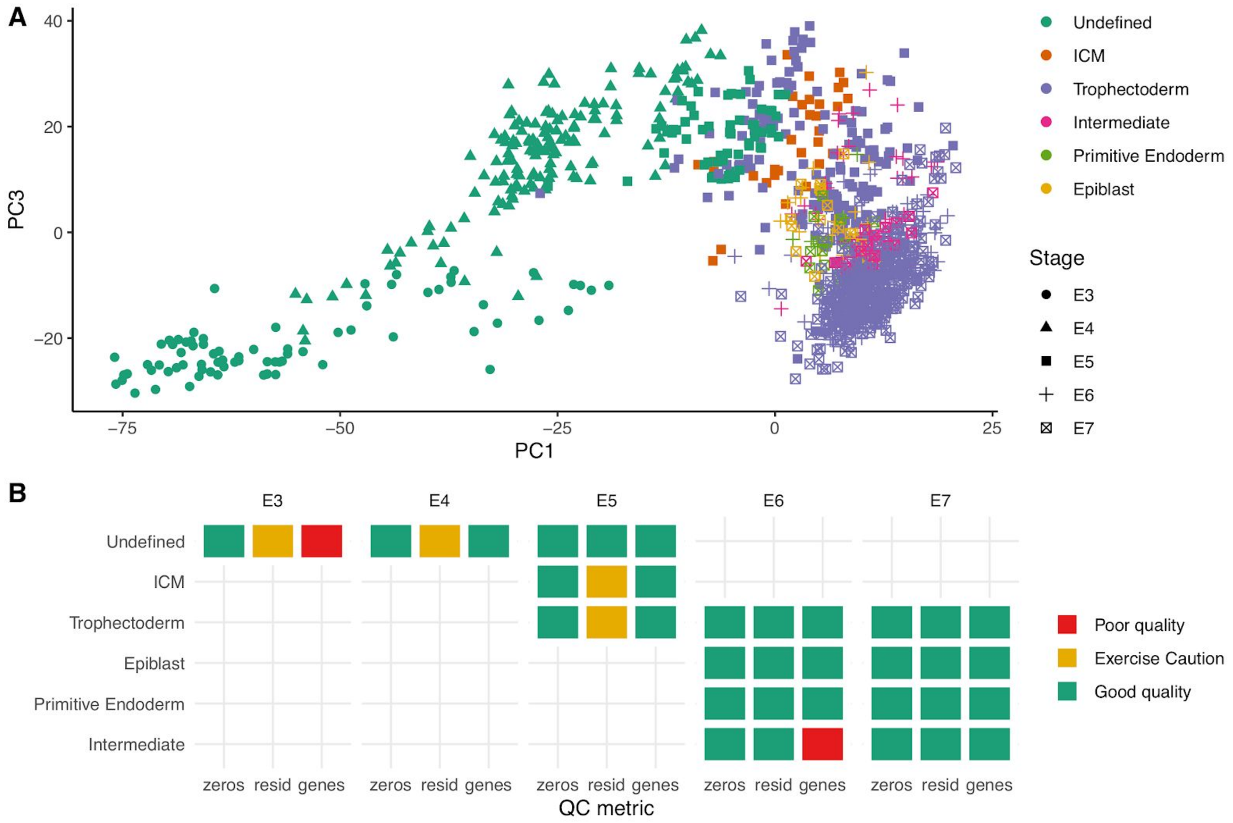


**Fig. S1.** Quality-control pre-processing of single-cell expression data. **A.** Principal component (PC) diagram of single-cell gene expression annotated with stage and cell type annotated based on Stirparo et al. (2018). These categories were used as grouping factors for *scploid*. PC1 and PC3 are depicted, as they best capture variation in stage and cell type, respectively. **B.** Quality control metrics computed by *scploid* for each stage/cell-type group. Groups failing a quality-control test (denoted as “Poor quality”) were excluded from subsequent analysis.

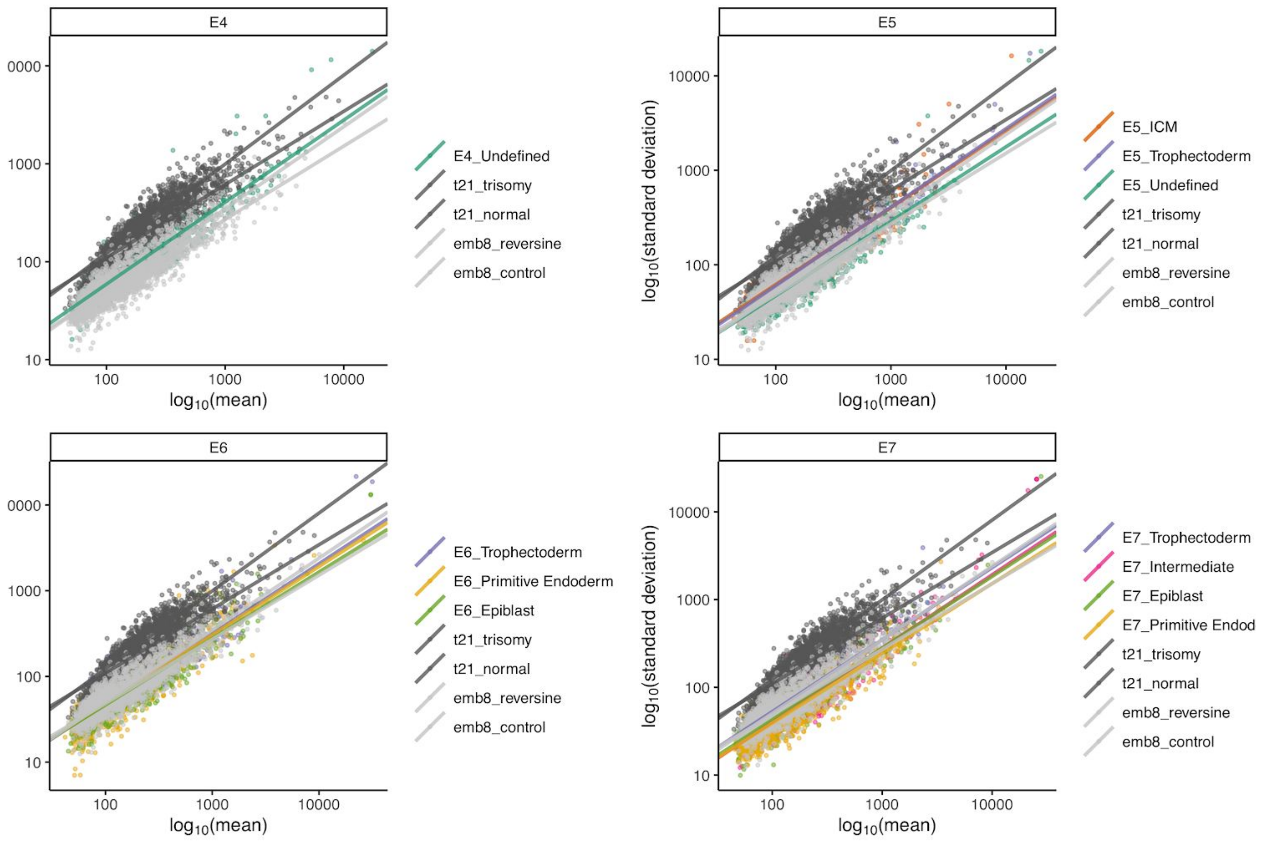


**Fig. S2.** Relationship between gene expression mean and standard deviation. Data from Petropoulos et al. (2016) are stratified by embryonic stage (days post-fertilization) and compared to mouse 8-cell embryo (emb8) and human trisomy 21 neuron (t21) data. These comparison data were previously used for *scploid* benchmarking by Griffiths et al. (2017) with good and poor performance, respectively.

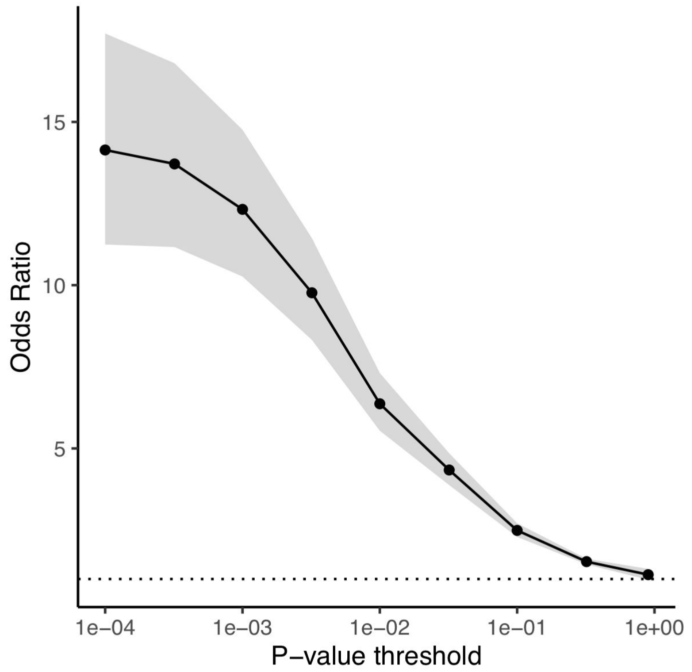


**Fig. S3.** Concordance between complementary signatures of aneuploidy. At increasingly stringent thresholds, we observe increasing overlap between chromosomes called as aneuploid based on evidence of expression alteration and allelic imbalance. Shaded region indicates 95% confidence interval.

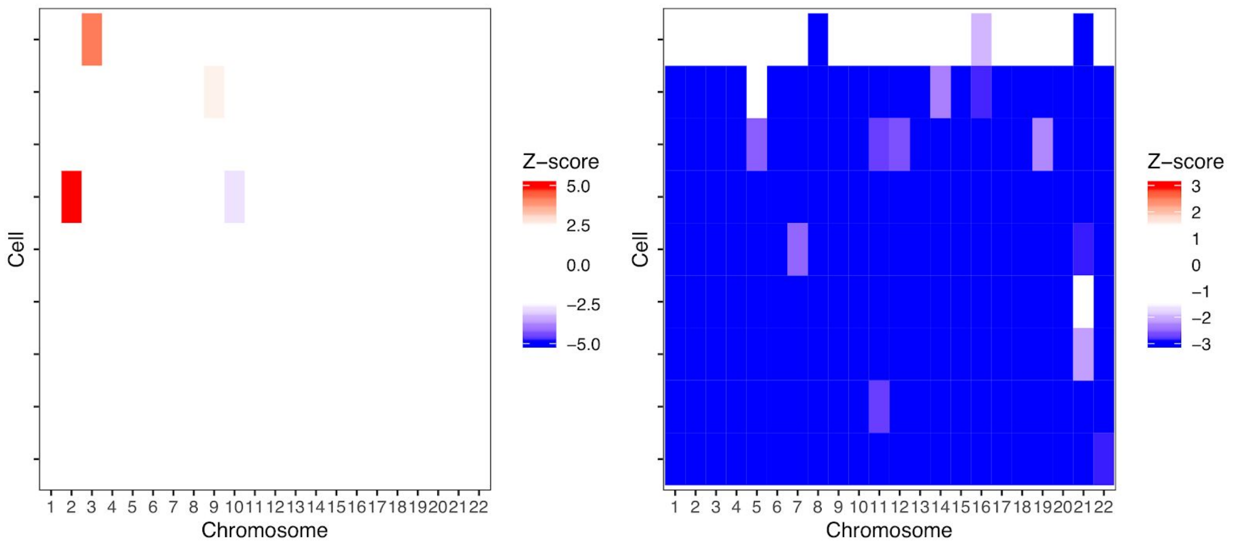


**Fig. S4.** Karyotype-wide abnormalities such as haploidy/near-haploidy are undetectable for embryo E7.5 based on expression signatures (left panel), but are evident based on widespread monoallelic expression (right panel). One cell (top row) exhibits mostly biallelic expression with only two monosomic chromosomes such that we classify the embryo as mosaic near-haploid.

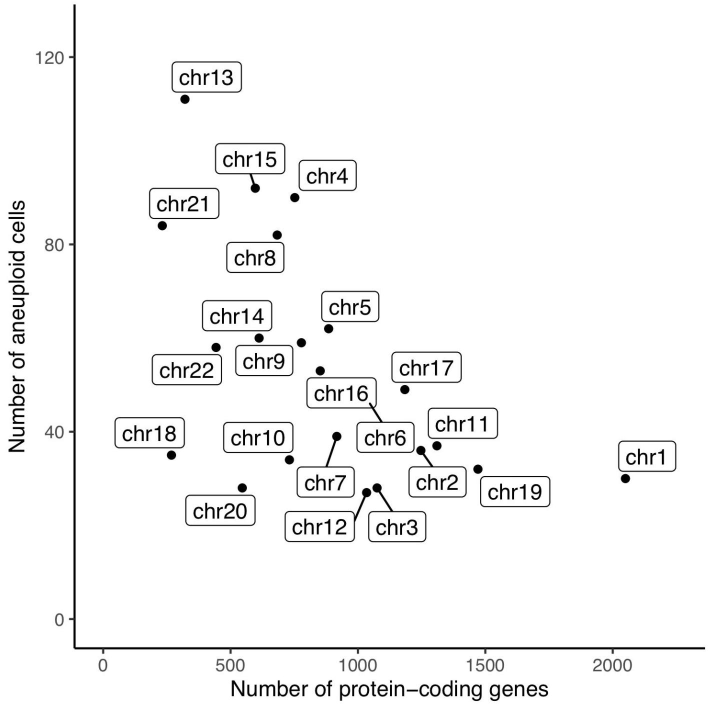


**Fig. S5.** Negative correlation between chromosome-specific aneuploidy rates and number of protein-coding genes per chromosome (Pearson's r = -0.546, $p = 8.64 \times 10^{-3}$). Differences in aneuploidy rates among chromosomes were not significant, however, after accounting for the correlation among chromosomes within cells within embryos ($\chi^2$(df = 21, n = 24,530) = 29.0, p = 0.114).

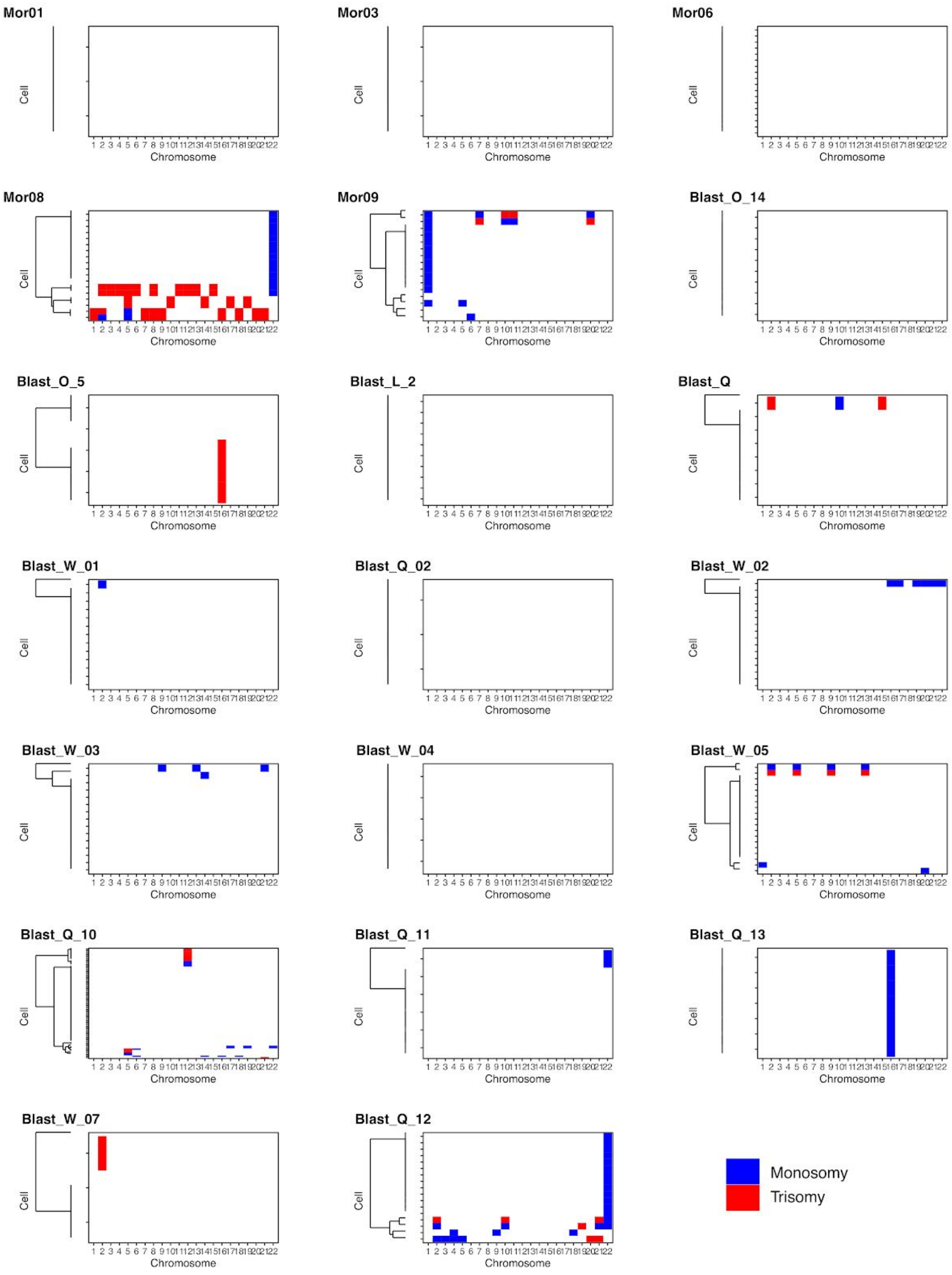


**Fig. S6.** Heatmaps of published aneuploidy calls based on PBAT scDNA-seq from Zhu et al. (2018b).

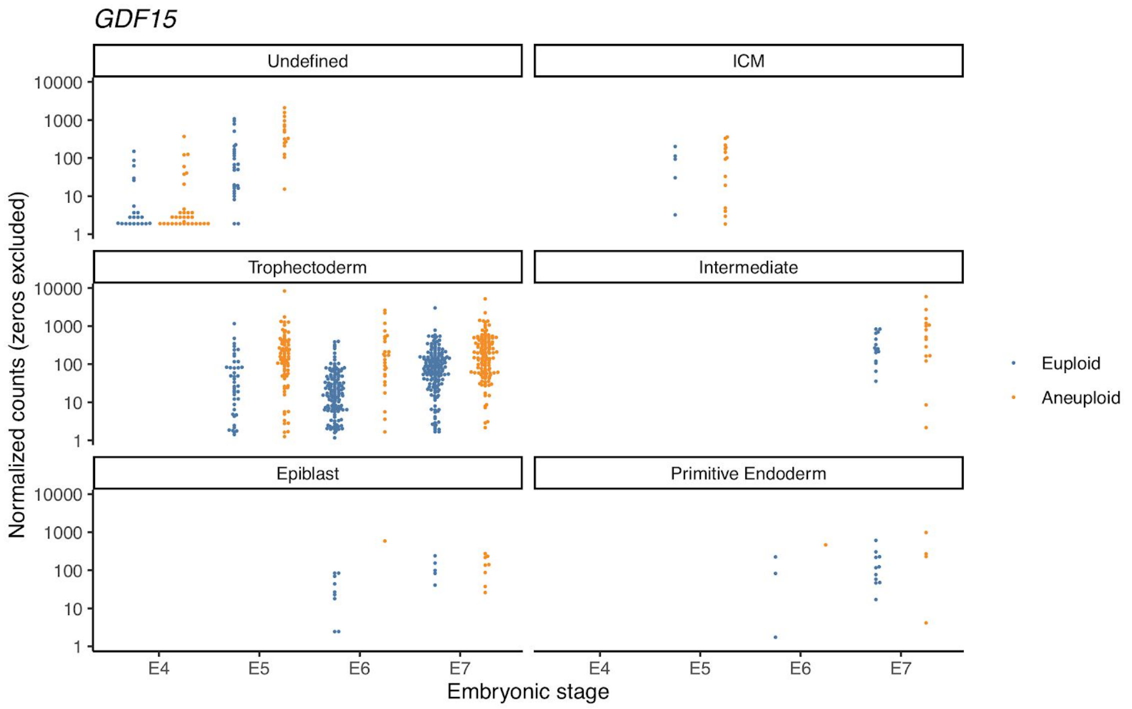


**Fig. S7.** Upregulation of *Growth/differentiation factor 15* (*GDF15*) in aneuploid compared to euploid cells.

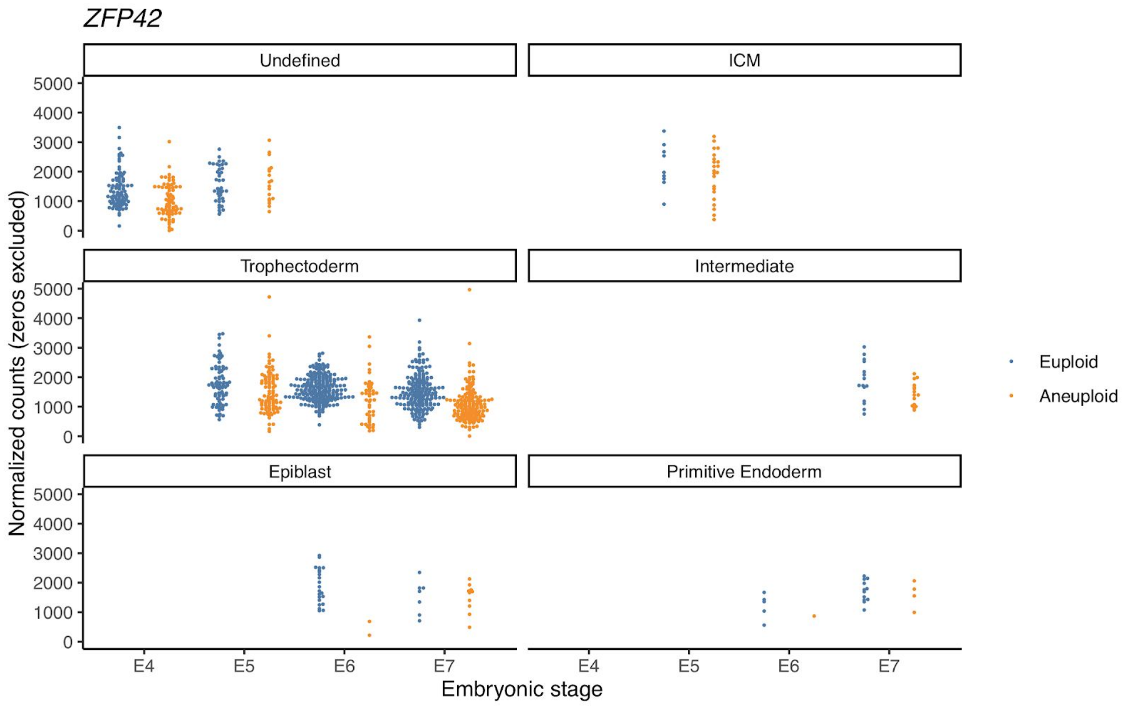


**Fig. S8.** Downregulation of *Zinc finger protein 42 homolog* (*ZFP42*) in aneuploid compared to euploid cells.

| | E4 | E5 | E6 | E7 | Total |
|---|---|---|---|---|---|
| **Embryos** | 16 | 23 | 18 | 17 | 74 |
| **Cells** | 160 | 262 | 290 | 403 | 1115 |
| Undefined | 160 | 60 | | | 220 |
| Inner cell mass | | 31 | | | 31 |
| Trophectoderm | | 171 | 260 | 340 | 771 |
| Intermediate | | | | 30 | 30 |
| Epiblast | | | 24 | 17 | 41 |
| Primitive endoderm | | | 6 | 16 | 22 |

**Table S1.** Counts and embryos and cells from Petropoulos et al. (2016) that were analyzed in this study, stratified by stage (days post-fertilization). Cells are further stratified by cell type.

| Symbol | β | SE (β) | P (β) | AME | SE (AME) | P (AME) |
|---|---|---|---|---|---|---|
| *GDF15* | 1.118 | 0.144 | $6.6 \times 10^{-15}$ | 142.06 | 58.40 | 0.015 |
| *ZFP42* | -0.262 | 0.037 | $1.6 \times 10^{-12}$ | -375.97 | 59.34 | $2.4 \times 10^{-10}$ |
| *POLR2E* | -0.214 | 0.031 | $6.3 \times 10^{-12}$ | -455.35 | 75.43 | $1.6 \times 10^{-9}$ |
| *PRPF31* | -0.314 | 0.047 | $2.6 \times 10^{-11}$ | -87.83 | 14.13 | $5.2 \times 10^{-10}$ |
| *NCL* | -0.156 | 0.024 | $9.1 \times 10^{-11}$ | -1102.83 | 174.51 | $2.6 \times 10^{-10}$ |
| *HNRNPC* | -0.168 | 0.027 | $2.4 \times 10^{-10}$ | -1170.02 | 189.51 | $6.7 \times 10^{-10}$ |
| *TUBA1B* | -0.254 | 0.041 | $4.9 \times 10^{-10}$ | -1587.99 | 331.07 | $1.6 \times 10^{-6}$ |
| *AHSA1* | -0.209 | 0.035 | $2.4 \times 10^{-9}$ | -328.67 | 58.29 | $1.7 \times 10^{-8}$ |
| *VDAC1* | -0.185 | 0.031 | $2.4 \times 10^{-9}$ | -345.92 | 74.55 | $3.5 \times 10^{-6}$ |
| *HMOX2* | -0.298 | 0.050 | $2.5 \times 10^{-9}$ | -100.93 | 19.75 | $3.2 \times 10^{-7}$ |
| *AAMP* | -0.236 | 0.040 | $3.4 \times 10^{-9}$ | -327.56 | 59.94 | $4.6 \times 10^{-8}$ |
| *AP000459.7* | 0.433 | 0.073 | $3.5 \times 10^{-9}$ | 1.85 | 0.39 | $2.1 \times 10^{-6}$ |
| *SEH1L* | -0.216 | 0.037 | $3.5 \times 10^{-9}$ | -178.47 | 30.49 | $4.8 \times 10^{-9}$ |
| *GPI* | -0.232 | 0.039 | $3.9 \times 10^{-9}$ | -344.82 | 76.69 | $6.9 \times 10^{-6}$ |
| *SLC4A1* | 0.487 | 0.084 | $5.5 \times 10^{-9}$ | 4.06 | 0.79 | $2.6 \times 10^{-7}$ |
| *SSH3* | -0.773 | 0.134 | $7.1 \times 10^{-9}$ | -20.73 | 5.80 | $3.4 \times 10^{-4}$ |
| *TBRG4* | -0.263 | 0.046 | $9.1 \times 10^{-9}$ | -222.93 | 41.54 | $8.0 \times 10^{-8}$ |
| *TRAP1* | -0.269 | 0.047 | $1.1 \times 10^{-8}$ | -239.05 | 52.04 | $4.4 \times 10^{-6}$ |
| *TIMM44* | -0.240 | 0.042 | $1.2 \times 10^{-8}$ | -159.01 | 29.99 | $1.1 \times 10^{-7}$ |
| *HARS* | -0.248 | 0.044 | $1.3 \times 10^{-8}$ | -190.60 | 36.85 | $2.3 \times 10^{-7}$ |

**Table S2.** Top twenty associations from analysis of differential expression expression comparing euploid and aneuploid cells. Regression coefficients (β) and average marginal effects (AME) are reported, along with corresponding standard errors and p-values. Positive coefficients indicate upregulation in aneuploid cells relative to euploid cells, while negative coefficients indicate downregulation.
